# NRPreTo: A Machine Learning Based Nuclear Receptor and Subfamily Prediction Tool

**DOI:** 10.1101/2022.11.12.516270

**Authors:** Sita Sirisha Madugula, Suman Pandey, Shreya Amalapurapu, Serdar Bozdag

## Abstract

The Nuclear Receptor (NR) superfamily includes phylogenetically related ligand-activated proteins, which play a key role in various cellular activities. NR proteins are subdivided into seven subfamilies based on their function, mechanism, and nature of the interacting ligand. Developing robust tools to identify NR could give insights into their functional relationships and involvement in disease pathways. Existing NR prediction tools only use a few types of sequence-based features and are tested on relatively similar independent datasets; thus, they may suffer from overfitting when extended to new genera of sequences. To address this problem, we developed Nuclear Receptor Prediction Tool (NRPreTo); a two-level NR prediction tool with a unique training approach where in addition to the sequence-based features used by existing NR prediction tools, six additional feature groups depicting various physiochemical, structural and evolutionary features of proteins were utilized. The first level of NRPreTo allows for the successful prediction of a query protein as NR or non-NR, and further subclassifies the protein into one of the seven NR subfamilies in the second level. We developed Random Forest classifiers to test on benchmark datasets, as well as the entire human protein datasets from RefSeq and Human Protein Reference Database (HPRD). We observed that using additional feature groups improved performance. We also observed that NRPreTo achieved high performance on the external datasets and predicted 59 novel NRs in the human proteome. The source code of NRPreTo is publicly available at https://github.com/bozdaglab/NRPreTo.

## Introduction

The Nuclear Receptor (NR) superfamily are transcription factors that are involved in critical signaling pathways controlling physiological growth, differentiation, and cell maintenance^1,2^. They respond to steroid-based signaling molecules like thyroid hormone, Vitamin D3, and retinoids. Due to their involvement in critical cell regulating activities, NRs are also instrumental in the disease pathways of several physiological and reproductive diseases such as breast cancer, obesity, and diabetes^3^. Structurally, all members of the NR superfamily have a common three-domain architecture. The N-terminal AF-1 region also known as the A/B region is a ligand-independent highly variable transcription activation region. They also contain two main domains; a moderately conserved Ligand Binding Domain (LBD) involved in ligand recognition and a highly conserved DNA Binding Domain (DBD) involved in DNA binding at the C-terminal region^4,5^. The DBD contains two conserved zinc finger motifs involved in Hormone Response Element (HRE) recognition^6,7^. Binding of the ligand at the LBD induces a conformational change in the DBD leading to the activation of specific HRE and subsequent downstream gene expression^8–10^. Hence both the conserved domains play important roles in NR-led gene regulation. The presence of both ligand and DNA binding domains in a protein sequence is therefore considered a signature hallmark of the NR superfamily. Since ligand binding is highly specific in the NR superfamily, these proteins are divided into seven subfamilies based on the type of ligand binding at the LBD. While NR1 to NR6 subfamilies are grouped clearly depending on their activating ligands, NR0 subfamily are characterized by “orphan receptors” with no known ligands and may sometimes lack DBD^11^. A detailed description of the subfamily names and their members can be found in Table S1.

NR is one of the most important superfamilies of druggable proteins. Some proteins of this superfamily are potential targets for developing therapeutic strategies for genetic diseases^12^. Furthermore, NR proteins are regulated by small molecule ligands making them amicable for drug designing strategies. These properties make NR promising pharmacological targets. Moreover, with the enormous number of protein sequences emerging from next generation sequencing efforts, there is a growing need to develop robust and accurate methods to identify NR and determine the subclass of the new incoming protein sequences.

In this regard, most existing tools for NR classification and subfamily prediction use only spectrum-kernel features, which are empirical in nature. One of the earliest NR prediction tools NRpred^13^, is a Support Vector Machine (SVM)-based tool that used only Amino Acid Composition (AAC) and Dipeptide Composition (DPC) features to classify proteins into four NR subfamilies. Gao et al.^14^ then improved NRpred by using Pseudo Amino Acid Composition (PAAC) features and achieved higher accuracy. However, both tools directly predict the NR subfamily before predicting if a given protein is NR or not. Later, NR-2L^15^ and iNR-PhysChem^16^ took a two-level prediction approach to address this problem and broadened the prediction scope to seven classes. At the first level, the tools determine if an incoming protein is an NR and at the second level, subfamily prediction is performed to classify the protein into one of the seven subclasses. This approach improved the overall accuracy but suffered from limitations like data redundancy and noisy features during model development. NRPred-FS^17^ addressed this issue through feature selection and achieved improved accuracy. Later, NRfamPred^18^ used the same two-level approach and utilized AAC and DPC features for level-1 and level-2 predictions, respectively. NRfamPred was verified on an independent dataset as well. Additionally they also carried out human proteome-wide NR subfamily prediction and identified 76 NR using the HPRD dataset^19^. However, NRfamPred did not apply any feature selection and used only the spectrum-kernel-like features for model building leading to lower overall performance. Recently, a Random Forest (RF)-based NR prediction tool RF-NR^20^ was developed using AAC, DPC, and Tripeptide Composition (TPC) features and achieved much higher performance. However, this tool has been tested only on independent datasets, which are relatively smaller, thus RF-NR may not be generalized enough to make proteome-scale NR predictions. Besides, RF-NR is not a publicly available tool.

In the current study, we hypothesized that since NRs differ from each other at allosteric sites^21^, merely using spectrum-like features to determine their subfamily cannot capture the structural differences among the proteins of different NR subfamilies. There is a need to consider additional features that account for evolutionary, co-evolutionary and physiochemical properties of proteins. Existing prediction tools use only primary sequence-based features to classify NR subfamilies. Here we investigated the role of various kinds of protein features that can govern NR subfamily classification. For this purpose, we developed Nuclear Receptor Prediction Tool (NRPreTo), which was trained on an extensive set of features to perform two-level NR prediction. Considering seven different feature groups including the three spectrum-kernel features (AAC, DPC and TPC) used by previous NR prediction tools, we computed 13,494 features. After performing feature selection, we trained RF models to predict whether an incoming protein is a NR or not and further predict its subfamily if found to be NR. We applied NRPreTo on two different data setups: (a) building a model using “train” dataset of a previously published benchmark dataset (BD) and testing on its independent dataset (b) combining the training and independent datasets of respective BD to generate a “combined” dataset used for training followed by testing on the human proteome collected from two external datasets; HPRD and NCBI RefSeq (humans)^22^. A schematic representation of the different components of NRPreTo is shown in Fig. 1.

**Figure 1.**
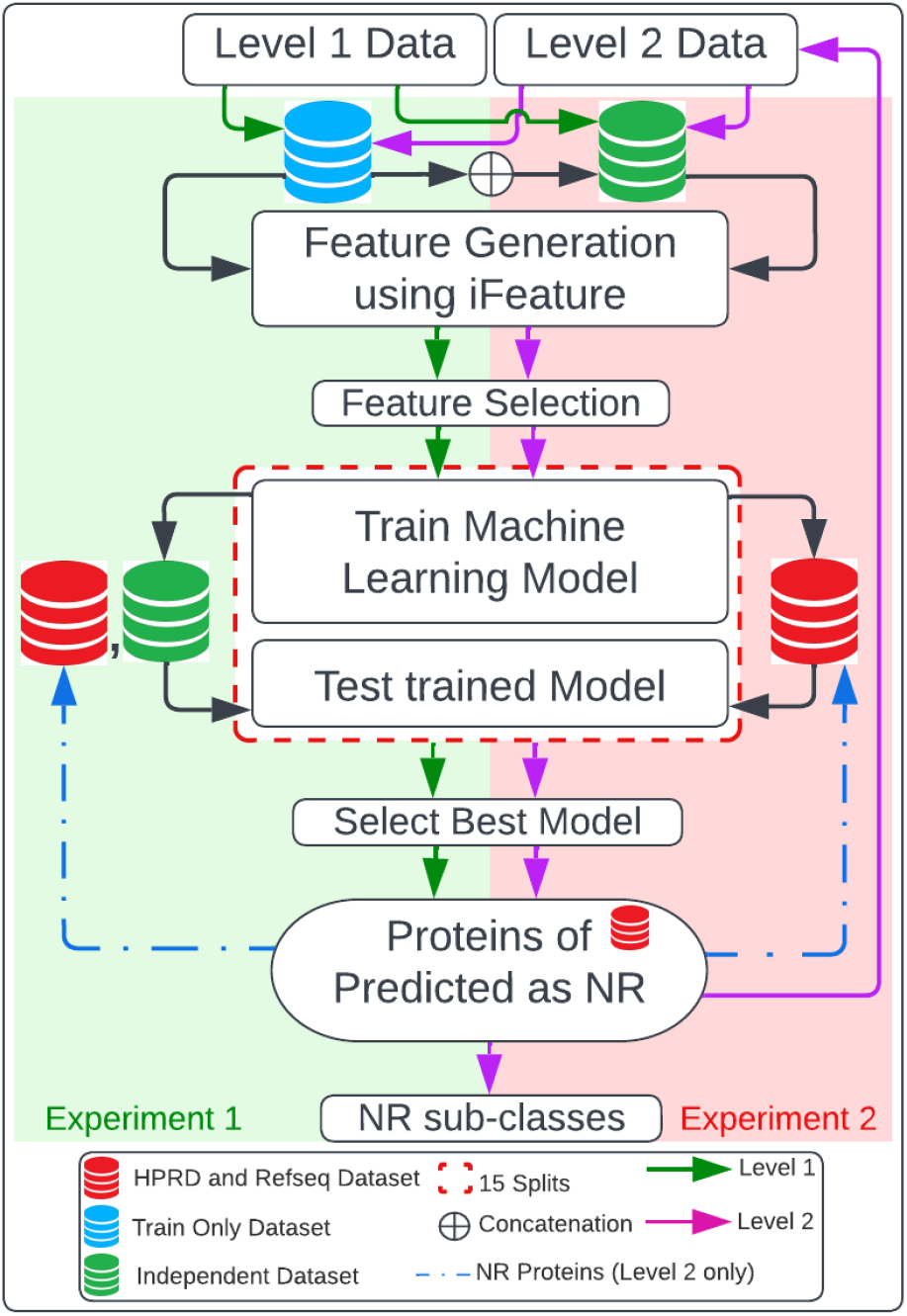
Overview of the NRPreTo workflow. Experiment 1 and 2 refers to testing the models on the independent datasets, and the human proteome dataset, respectively. Level 1 is the prediction task to classify NR proteins and level 2 is the prediction task to classify the subfamily of an NR protein.

Our proposed methodology has outperformed or been on par with all previous studies. Additionally, we also predicted 59 new NRs from the human proteome. Some of the identified proteins include pharmacologically important drug targets, which can be promising candidates for further studies. To the best of our knowledge, this is the first NR classification study to build classifiers using the entire spectrum of protein features in addition to the regularly used sequence-based features. Moreover, this is the first study to test its models not only on two independent datasets but also carry out a proteome scale prediction across two important public datasets: HPRD and RefSeq covering all sequences of human proteome known so far. These results indicate that NR classification is influenced by other protein features in addition to sequence based features.

## Results

In the current study we developed a two-level NR prediction tool called NRPreTo, built on a multitude of protein features (Table 1) in addition to the sequence-based features used by the previous studies. Using two experimental setups with different sized training datasets (see Materials & Methods), we performed extensive evaluations of NRPreTo on two previously published BDs (i.e., BD1 and BD2) and the entire human proteome. We successfully identified 59 new NRs from human proteome while improving the performance of the model.

**Table 1.** List of descriptors used in the study

| **Table 1** List of descriptors used in the study |  |  |
| --- | --- | --- |
| **Descriptor Group** | **Descriptor set** | **Descriptors** |
| Amino Acid Composition | Amino acid composition (AAC) | 20 |
|  | Composition of k-spaced amino acid pairs (CKSAAP) | 2400 |
|  | Dipeptide composition (DPC) | 400 |
|  | Dipeptide deviation from expected mean (DDE) | 400 |
|  | Tripeptide composition (TPC) | 8000 |
| Grouped Amino Acid Composition | Grouped amino acid composition (GAAC) | 5 |
|  | Composition of k-spaced amino acid group pairs (CKSAAGP) | 150 |
|  | Grouped dipeptide composition (GDPC) | 25 |
|  | Grouped tripeptide composition (GTPC) | 125 |
| Autocorrelation | Moran (Moran) | 240 |
|  | Geary (Geary) | 240 |
|  | Normalized Moreau-Broto (NMBroto) | 240 |
| C/T/D | Composition (CTDC) | 39 |
|  | Transition (CTDT) | 39 |
|  | Distribution (CTDD) | 195 |
| Conjoint Triad | Conjoint triad (CTriad) | 343 |
|  | Conjoint *k*-spaced triad (KSCTriad) | 343 |
| Quasi-Sequence-Order | Sequence-order-coupling number (SOCNumber) | 60 |
|  | Quasi-sequence-order descriptors (QSOrder) | 100 |
| Pseudo-Amino-Acid Composition | Pseudo-amino-acid composition (PAAC) | 50 |
|  | Amphiphilic PAAC (APAAC) | 80 |

\*Table adopted from iFeature package

### Level-1 predictions

We carried out level-1 prediction to screen-in query proteins that are NRs. Only the sequences determined as NR using the level-1 models were taken ahead to the second level for subfamily classification. Since this level is a binary classification problem, both NR and Non-NR sequences were used to train the models.

We used BorutaPy feature selection method^23^ to select separate important features sets for level-1 models for the two experimental setups. Fig. 2 shows a feature-set-wise summary of selected features of level-1 model for experiment-1 (“train” data) and experiment-2 (“combined” dataset). For experiment-1, 557 and 385 important features were selected for BD1 and BD2, respectively, whereas for experiment-2, 2635 and 702 important features were selected for the two BD, respectively. In both BDs, the maximum number of important descriptors were from Amino acid composition group followed by C/T/D (Fig. 2). Amino acid composition group in this study include CKSAAP, TPC and DDE descriptor sets, which constitute the maximum count of their group (see Table 1). These are sequence-based features, which are elemental to any protein and are widely employed in different protein identification and classification tasks^24–30^. Additionally, other descriptor sets like PAAC (Pseudo Amino Acid Composition), CKSAAP (Amino Acid Composition) and Moran (Autocorrelation) were also identified among the important features, which are also known to play significant role in classification of proteins, including NR^31–35^. PAAC, which represents positional and compositional pattern of amino acid in protein sequences preserve the evolutionary information of the proteins. Therefore, it is widely used in problems like predicting various post-translational modification sites or identifying protein subcellular localization where evolutional relationships of residues are important ^36–41^.

**Figure 2.**
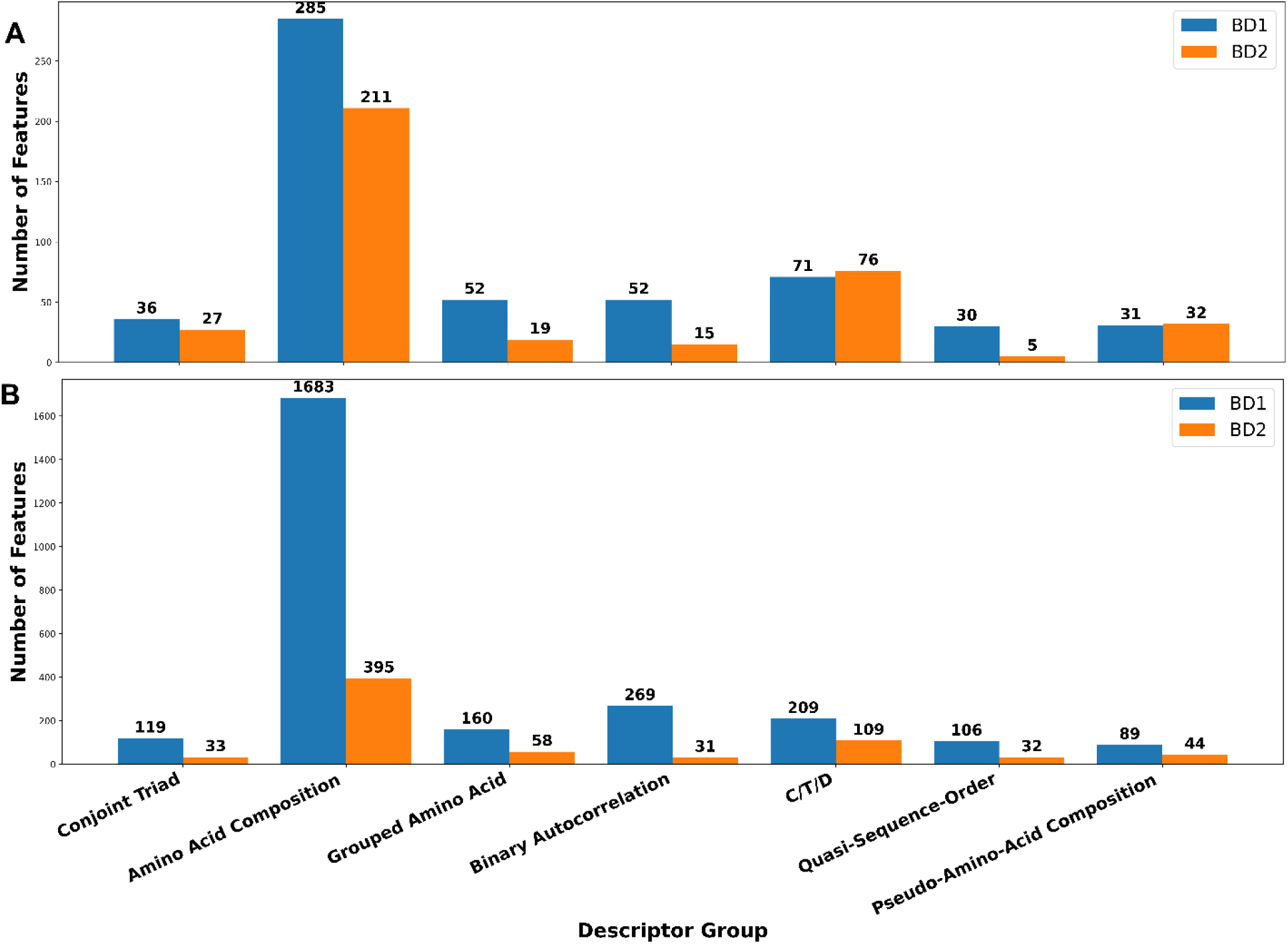
Feature set-wise distribution of the important descriptors of the level-1 prediction model. **A)** Experiment-1 (train only data) and **B)** Experiment-2 (combined dataset).

#### Experiment-1 – Train Only

Experimental setup-1 consists of models trained only on the “training” dataset of the two BDs and tested on their respective independent datasets. The BD1 training contained 267 NR and 1000 non-NR sequences while BD2 had 159 NR and 500 non-NR (Table S1). Due to the imbalance in the dataset, class weights were used to give equal importance to the two classes and negate the influence of the majority class during prediction. A comparison of the average performance of train only models on independent datasets over 15 runs across the two BDs is shown in Table 2. Further, we compared NRPreTo with four existing NR prediction tools namely NR-2L, NRPred-FS, NRfamPred and RF-NR to assess its performance. Since we used the same BD that were used to develop the aforementioned tools, we did a one-to-one comparison of NRPreTo’s best model with the results reported by the existing tools. We selected the best model for NRPreTo based on F1 and AUC scores. We also reported the performance of our method over 15 runs to observe variance of the model. On BD1, we compared our model performance with NRPred-FS and RF-NR, which were trained on BD1 while for BD2 we compared our results with NR-2L, NRfamPred and RF-NR, which were trained on BD2.

**Table 2.**
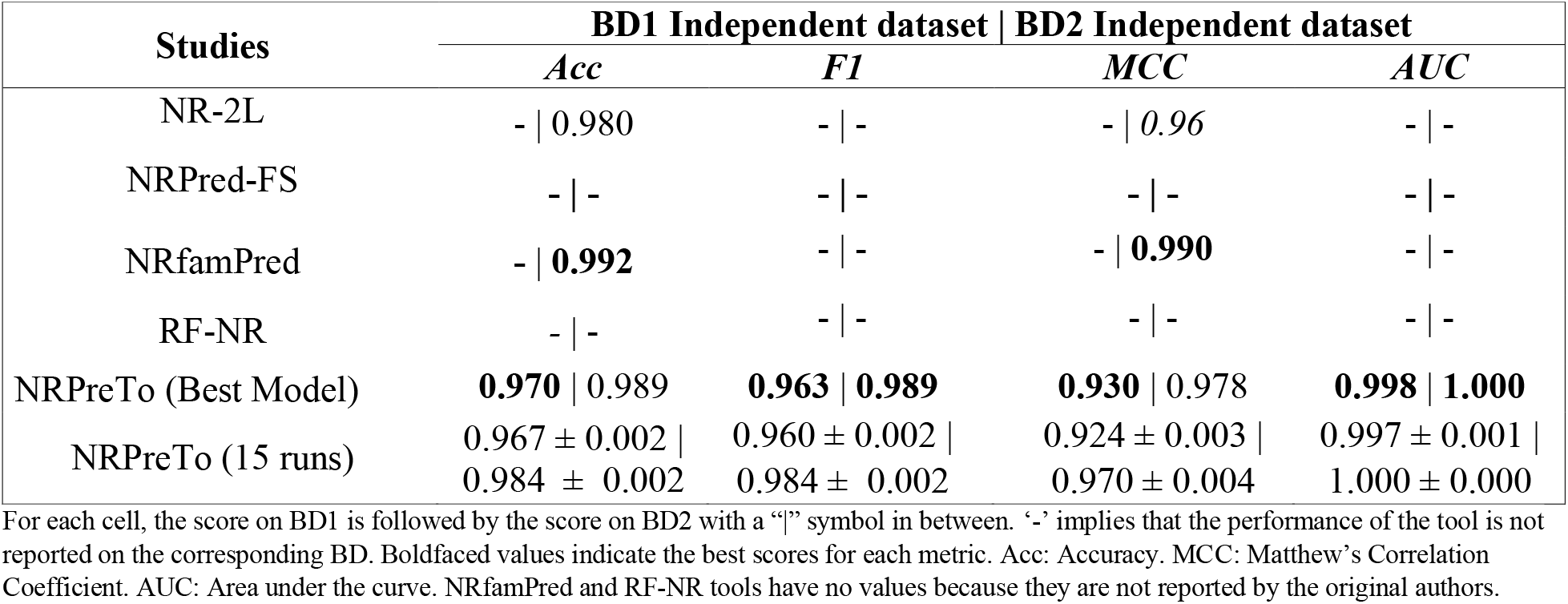
Performance comparison of NRPreTo level-1 “Train only” models with existing tools on the two independent datasets.

Due to the several missing values in the papers of the previous tools, a comprehensive comparison was not possible. The source codes of the tools were not available to test directly, either. Based on the available results, however, we observed that NRPreTo’s performance was on par with the performance of the other tools (Table 2).

#### Experiment-2 (Combined Dataset)

To test the performance of NRPreTo on the entire human proteome, we trained a new RF model using a “combined dataset” of training and independent sets for each BD. The combined BD1 dataset contained 3016 NR and 2064 Non-NR proteins while combined BD2 consisted of 727 NR and 1000 Non-NR proteins, respectively. A summary of the prediction performance of RF models on HPRD and RefSeq datasets over 15 runs using the combined models is given in Table 3.

**Table 3.**
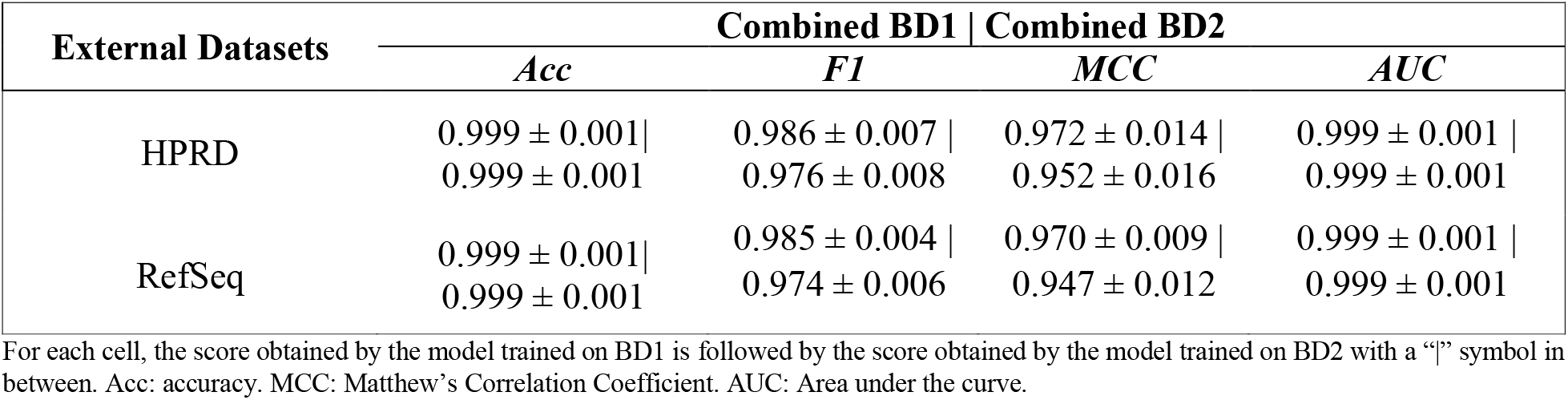
Summary of level-1 prediction performance of combined models on the HPRD and RefSeq datasets over 15 runs.

| External Datasets | Combined BD1 \| Combined BD2 |  |  |  |
| --- | --- | --- | --- | --- |
|  | *Acc* | *F1* | *MCC* | *AUC* |
| HPRD | 0.999 ± 0.001 \| | 0.986 ± 0.007 \| | 0.972 ± 0.014 \| | 0.999 ± 0.001 \| |
|  | 0.999 ± 0.001 | 0.976 ± 0.008 | 0.952 ± 0.016 | 0.999 ± 0.001 |
| RefSeq | 0.999 ± 0.001 \| | 0.985 ± 0.004 \| | 0.970 ± 0.009 \| | 0.999 ± 0.001 \| |
|  | 0.999 ± 0.001 | 0.974 ± 0.006 | 0.947 ± 0.012 | 0.999 ± 0.001 |

For each cell, the score obtained by the model trained on BD1 is followed by the score obtained by the model trained on BD2 with a “|” symbol in between. Acc: accuracy. MCC: Matthew’s Correlation Coefficient. AUC: Area under the curve.

Our “BD1 combined dataset” RF model successfully predicted 246 NRs out of 249 NRs in RefSeq dataset and all 127 NRs from the HPRD dataset. On the other hand, “BD2 combined dataset” RF model was able to correctly classify 234 and 122 NRs in the RefSeq and the HPRD datasets, respectively. Among the existing NR prediction tools, only NRfamPred carried out proteome scale prediction. They used only HPRD dataset and utilized limited number of features for model generation, which does not account for the different structural, functional profiles of the NR proteins. Besides, NRfamPred identified only 76 novel NRs across HPRD dataset. It has been demonstrated that using a combination of different protein features along with appropriate feature selection methods can improve prediction performance of classification models^42^. In line with this, our tool was able to achieve higher prediction performance in terms of accurately identifying maximum number of NRs across human proteome reported so far by any previous studies or databases.

### Level-2 Predictions

In level-2 prediction, the goal was to determine the subfamily of an NR protein. Only the proteins that were predicted as NR at level-1 were taken ahead for level-2 classification. We trained RF models using only the NR sequences belonging to the seven NR subfamilies considered in the study. We used a similar feature selection process in selecting important features for level-2 as in level-1. For experiment-1, 402 and 407 important features were selected for BD1 and BD2, respectively, whereas for experiment-2, 2870 and 835 important features were selected for BD1 and BD2, respectively (Fig. 3). Further, amino acid composition group constituted the maximum number of important features in level-2 as in level-1.

**Figure 3.**
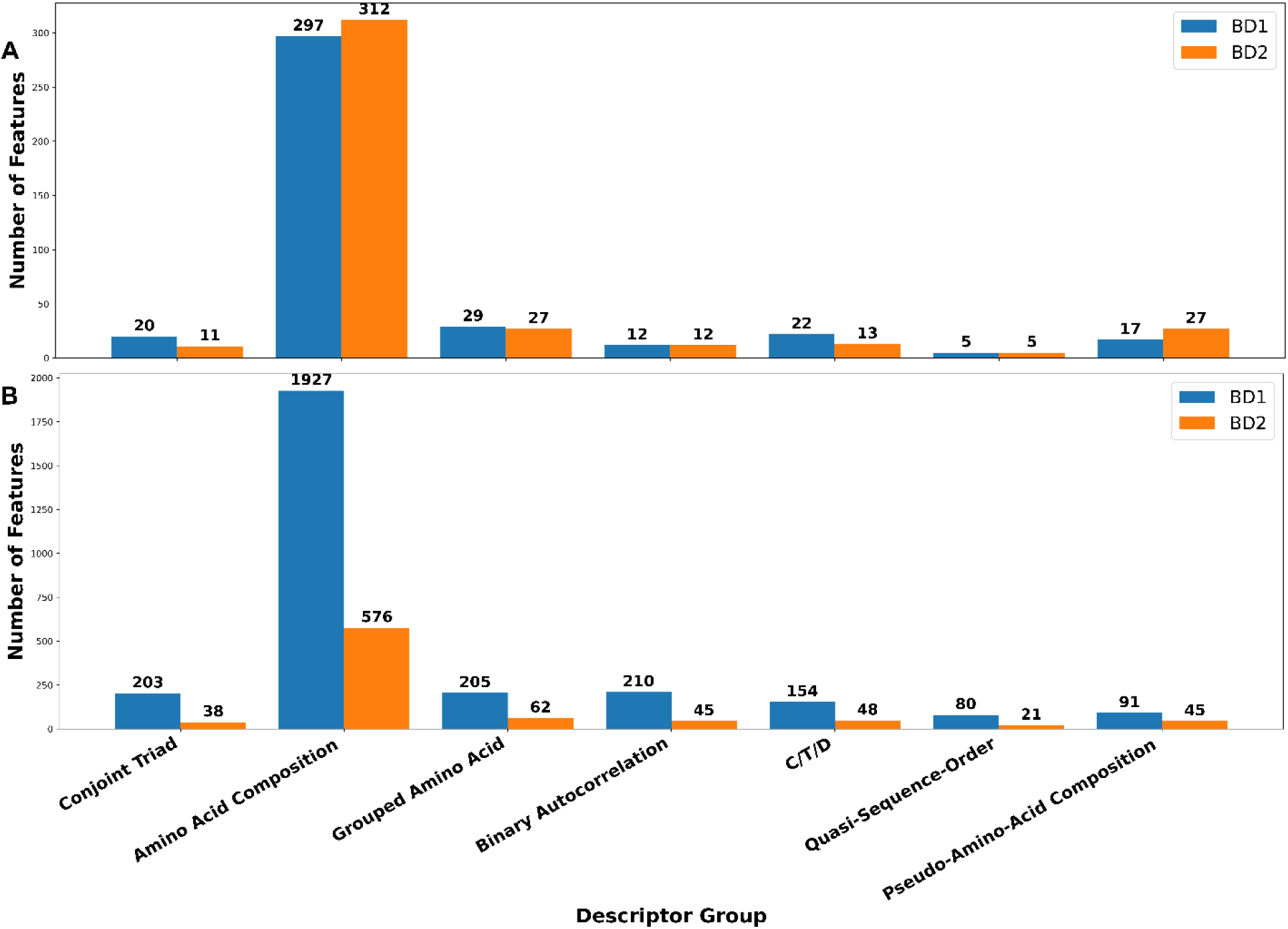
Feature set-wise distribution of the important descriptors of level-2 prediction model. **A)** Experiment-1 (train only) and **B)** Experiment-2 (combined dataset).

#### Experiment-1 (Train Only)

As in experimental setup 1 of level 1, this dataset also contains only the training set sequences (i.e., without including independent set). However, since level-2 is a multiclass classification task, the classifiers were trained only on the positive set (i.e., NR sequences) to classify an incoming protein into one of the seven NR subfamilies. Thus, the training dataset in this experiment consisted of 267 and 159 NRs for BD1 and BD2, respectively. A breakup of the number of sequences in every subfamily class in BD1 and BD2 is given in Table S1. As seen in Table S1, the number of training sequences in some classes were fewer compared to the others leading to a highly imbalanced dataset. Therefore, class weights were used as in level-1 to give equal weight to all the seven classes during model training.

We compared the level-2 prediction performance of NRPreTo with the existing tools. As seen in Table 4, most of the existing tools are built only on one of the BDs and tested on only one of the independent datasets. Also, these tools are reported using only few evaluation metrices on the independent dataset. Although RF-NR used both BDs in their pipeline, they reported the model performance using only MCC and recall for BD2 independent dataset. NRPreTo is the first tool to build models using both BDs and evaluate them on their respective independent datasets. With the evaluation profiles of the existing tools, it is difficult to derive conclusions about the overall model performance. Based on the available values, the best NRPredTo model had almost comparable accuracy to RF-NR on BD1, but outperformed it on both BDs based on MCC and F1 scores, which are reliable metrices for imbalanced classes (Table 4). It also outperformed NRfamPred and NR-2L on BD2 based on accuracy and MCC, respectively.

**Table 4.** Performance comparison of NRPreTo level-2 RF “Train only” models with existing tools on the two independent datasets.

|  |  |  |  |  |
| --- | --- | --- | --- | --- |
| **Studies** | **BD1 Independent dataset \| BD2 Independent dataset** |  |  |  |
|  | *Acc* | *F1* | *MCC* | *AUC* |
| NR-2L | - \| 0.996 | - \| - | - \| - | - \| - |
| NRPred-FS | 0.974* \| - | - \| - | - \| - | - \| - |
| NrfamPred | - \| 0.995 | - \| - | - \| 0.980 | - \| - |
| RF-NR | **0.996** \| - | 0.989 \| - | 0.980 \| 0.990 | - \| - |
| NRPreTo<br>(Best Model) | 0.992 \| **0.998** | **0.998** \| **0.999** | **0.987** \| **0.991** | **0.989** \| 0.997 |
| NRPreTo<br>(15 runs) | 0.983 ± 0.004 \|<br>0.993 ± 0.003 | 0.997 ± 0.000 \|<br>0.998 ± 0.000 | 0.957 ± 0.009 \|<br>0.967 ± 0.015 | 0.977 ± 0.005 \|<br>0.991 ± 0.004 |

For each cell, the score on BD1 is followed by the score on BD2 with a “|” symbol in between. \* Indicates that the accuracy value reported for NRPred-FS is taken from the RF-NR publication. Since NRPred-FS was not evaluated on BD1 independent dataset, this value is taken from RF-NR publication who calculated NRPred-FS accuracy on BD1 independent set. Boldfaced values indicate the best scores for each metric. Acc: accuracy. MCC: Matthew’s Correlation Coefficient. AUC: Area under the curve.

#### Experiment-2 (Combined Dataset)

To perform level-2 classification on the human proteome, we trained models on the combined datasets and tested these models on the human proteome. Here the classifiers of BD1 and BD2 were trained on 3016 and 727 NR sequences, which makeup the number of sequences in the seven NR subfamilies. The class imbalance issue was address using class weights as in the previous experiments.

The results indicated that the models produced consistent results across both the external datasets over the 15 splits (Table 5). The combined BD2 model correctly classified the subfamilies of 120 out of 122 NRs identified at level-1 from HPRD. It also classified subfamilies of 230 RefSeq sequences correctly out of 234 NRs identified at level 1. The model trained on combined BD1 on the other end correctly classified 125 out of 127 HPRD NRs and 241 out of 246 RefSeq NRs identified at level 1. These results indicate that the descriptors chosen at this level were highly contributing and produced reliable results regardless of the number of splits.

**Table 5.** Summary of level-2 RF models prediction performance trained on combined models BD1 or BD2 dataset and tested on external datasets over 15 runs.

|  |  |  |  |  |
| --- | --- | --- | --- | --- |
| **External Datasets** | **Combined BD1 \| Combined BD2** |  |  |  |
|  | *Acc* | *F1* | *MCC* | *AUC* |
| HPRD | 0.9843 ± 0.0084 \| | 0.9423 ± 0.0018 \| | 0.9773 ± 0.0001 \| | 0.9764 ± 0.00079 \| |
| | 0.9836 ± 0.0270 | 0.9423 ± 0.0124 | 0.9763 ± 0.0027 | 0.9920 ± 0.0039 |
| RefSeq | 0.9786 ± 0.0018 \| | 0.9595 ± 0.0007 \| | 0.9676 ± 0.0027 \| | 0.9745 ± 0.0058 \| |
| | 0.9829 ± 0.0068 | 0.9620 ± 0.0245 | 0.9741 ± 0.0150 | 0.9740 ± 0.0052 |

For each cell, the score obtained by the model trained on BD1 is followed by the score obtained by the model trained on BD2 with a “|” symbol in between. Acc: accuracy. MCC: Matthew’s Correlation Coefficient. AUC: Area under the curve.

### Analysis of the importance of protein feature categories

We performed feature importance analysis for the models at both levels to assess the influence of different descriptor groups on the performance of our models. For this, we performed SHapley Additive exPlanations (SHAP) analysis^43^, which utilizes a game theoretic approach^44^ to compute the contribution of each feature in predictions. Since our best performing models were on the combined datasets, we chose to do a feature importance analysis of RF models trained on the combined datasets. For this purpose, we used the best model of the combined dataset (Table 3 and 5) and computed the average SHAP value for each of the seven descriptor groups identified during feature selection.

The SHAP analysis results showed that for all models, the features from all descriptor groups had contribution (Fig. 4). Particularly, features from the amino acid composition group namely AAC, DPC, TPC, Composition of K-spaced Amino Acid Composition (CKSAAP) and Dipeptide Deviation from Expected mean (DDE) had the highest mean SHAP value. This trend agrees with the fact that this group of descriptors consistently formed the highest fraction of the important features on both of the combined models. The previous studies utilized features from “Amino Acid Composition” group, too. However, they only considered AAC, DPC and TPC feature sets. Our results indicate that features from CKSAAP and DDE were the most contributing for two models (Fig. 4A and 4D) and second most contributing in the other two models (Fig. 4B and 4C). It is known that CKSAAP features can capture patterns of short linear motif information and thereby preserve the coevolutionary information of amino acid residues because of which they are employed in post-translational modification site identification^45–47^.

**Figure 4.**
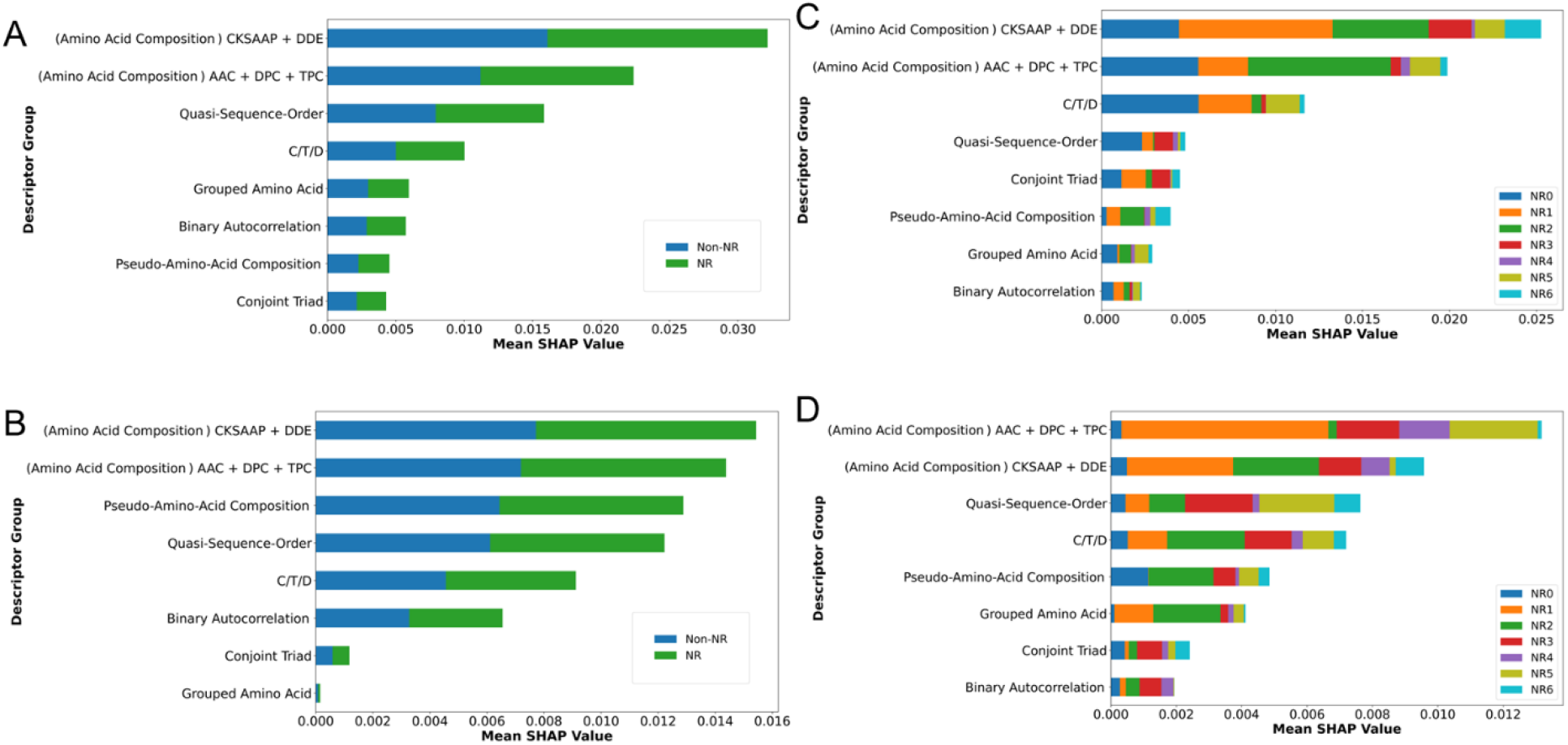
Feature importance analysis. SHAP analysis was performed for each feature and mean SHAP value for each description group was calculated for **(a)** level 1 model using BD1 **(b)** level 1 model using BD2 **(c)** level 2 model using BD1 and **(d)** level 2 model using BD2. See Table 1 for the acronyms of the descriptor groups.

### NRPreTo predicts known and novel NRs

NRPreTo classified a total of 253 sequences from both external sets. We noticed that among the 76 sequences reported by NRfamPred, our tool classified 64 sequences as NR and the remaining 12 sequences were predicted as non-NR. Upon careful analysis of these 12 sequences in NCBI-PROTEIN database, we did not observe the signature ligand and DNA binding domains of NR. Moreover, UniProt and HPRD do not annotate these sequences as NRs (Table S2). Thus, these sequences are unlikely to be a NR. There were 61 sequences whose subfamily class is reported in HPRD and UniProt but not by NRfamPred. NRPreTo was able to predict the subfamily of all these sequences correctly (Table S3). NRPreTo also correctly predicted 60 sequences, which are not annotated as NR in HPRD, has no reported NRfamPred prediction, but UniProt annotated them as NR (Table S4). We also predicted the subclasses of 59 RefSeq sequences, which are not annotated as NR in HPRD and UniProt (Table S5). Upon further analysis of these sequences in NCBI-PROTEIN database, we noticed that all 59 sequences have the signature conserved ligand and DNA binding domains of a classic NR. Therefore, we propose these 59 sequences as the novel predictions of NRPreTo.

## Discussion

In the current study, we developed an NR prediction tool called NRPreTo, which performs two-level classification to identify NRs and their subfamilies. We evaluated NRPreTo on two BDs published in earlier studies and on human proteome from HPRD and RefSeq datasets. Compared to the existing NR prediction tools, which are built only using some sequence-based features, NRPreTo was developed using a complete spectrum of protein features including the sequence-based features. We made NRPreTo publicly available at https://github.com/bozdaglab/NRPreTo under Creative Commons Attribution Non-Commercial 4.0 International Public License.

This study was carried out at under two experimental setups. In the first setup, we built a model using the training datasets of each of the BDs, and testing on the corresponding independent dataset in the BD. In the second setup, we trained a model using both training and independent datasets of each of the BDs and tested on the human proteome from HPRD and RefSeq. We observed that the performance of the models (developed using different protein features) on independent datasets was on par with the existing tools. Moreover, the strategy of combining the datasets to train the classifiers further boosted the performance. While comparing Table 3 with Table S6 and Table 5 with Table S7, it is seen that the models trained on combined datasets achieved much higher performance than the models trained using the training data only. For level 2, the performance of combined models was higher than train-only models for all metrics except for F1 on HPRD dataset (Table S7). Using this strategy, we not only predicted subfamilies of 60 sequences unannotated by HPRD but also proposed 59 novel NR sequences that are not annotated as NR in any databases. Using the feature importance analysis on the combined models, we observed features from CKSAAP and DDE descriptor groups contributed highly to the prediction tasks at both levels (Fig. 4).

Our comparisons with existing tools were limited by the lack of scores for some evaluation metrics reported in their respective papers. All the existing tools had models trained and tested using only one of the BDs. They did not have the source code available for us to run their models. Moreover, except NRfamPred, no other tool has tested their model to identify new NRs from the human proteome. Thus, we could not compare NRPredTo performance with all tools on all BDs for all performance metrics.

## Materials and Methods

### Data Sources

#### Benchmark Datasets

We utilized two previously published benchmark datasets, namely benchmark dataset 1 (BD1) of RF-NR and benchmark dataset 2 (BD2) of NR-2L. These BDs are composed of training and independent sets containing both NR and Non-NR sequences (Table S1). Both BDs were prepared from the Nuclear Receptor Database (NucleaRDB Release5.0) containing 3016 NR sequences, which are phylogenetically classified into seven subfamilies with each subfamily containing NR sequences from different animal species^48^. The detailed procedure of generating the BDs is outlined in NR-2L^15^ and RF-NR^20^ studies.

#### External Datasets

To further evaluate the models trained on BD, we used two high quality human proteome datasets, namely HPRD and NCBI-RefSeq. HPRD dataset contains 30,046 manually curated human proteins. Since HPRD dataset could not be downloaded from HPRD website, we used the data distributed by OmniPath rescued data repository^49^ for this study. We verified that the data obtained from OmniPath is the latest version of HPRD (HPRD_Release 9.0) mentioned on the HPRD website. Since the version on HPRD website is not updated after 2009, we also tested our models on another updated set of human protein sequences derived from NCBI-RefSeq. NCBI-RefSeq database contains integrated, comprehensive, non-redundant, well-annotated sequences of genomic DNA, transcripts and proteins for various organisms. For this study, we only considered human protein sequences reported in RefSeq database, which had 116,585 records at the time of download. After eliminating the computationally predicted models (i.e., the records with accession numbers prefixed with “XP” or “YP”), we had 62,625 records, which were used in this study.

### Protein Feature Generation and Feature Selection

For descriptor calculation, we used iFeature^50^, an open-source Python package to compute a multitude of protein and peptide features from their amino acid sequences. The features included structural, physiological, evolutionary, and pattern-based descriptors depicting the complete profile of a protein. Although the program offers 18 descriptor groups, we calculated descriptors belonging to only seven groups for this study as these are the only groups that could be computed using the FASTA sequences of different length. Each group contains multiple descriptors sets and each descriptor set contains multiple descriptors. The seven groups considered in this study contained 21 descriptor sets that made up of 13,494 descriptors (Table 1).We applied Borutapy feature selection method to identify the important features governing NR classification. Here level-1 and level-2 were considered as two separate classification problems and feature selection was done separately on both levels.

### NRPreTo Experimental Settings

To evaluate NRPreTo, we conducted two experiments. In the first experiment, a model was trained using the “training data” of BD1/BD2 and tested on the corresponding “independent data”. In the second experiment, models were built on a “combined dataset” of training and independent sets for the respective BD1/BD2 and tested on the external datasets. Random Forest (RF) was employed for model building.

We used a 7-fold CV for hyperparameter tuning using Hyperopt^51^, a Python package that utilizes a Bayesian optimization technique to find best hyperparameter combination capable of producing the best fit. We tuned five hyperparameters (Table S8). To combat the class imbalance in the training dataset, we assigned weights to the individual classes. The class weights were inversely proportional to the class frequency. This ensured that the model gives equal importance to every class and therefore pushes the models to learn a better representation of the minority class. To observe the variance of the model, we trained and tested our model 15 times. We used several evaluation metrics such as F1 score, area under the ROC curve (AUC-ROC) to validate our models. Then an average and a standard deviation of the model’s performance on the independent or external datasets across the 15 runs were reported.

## Acknowledgements

This work was supported by the National Institute of General Medical Sciences of the National Institutes of Health under Award Number R35GM133657.

## References

1. Mangelsdorf, D. J. et al. The nuclear receptor superfamily: The second decade. Cell 83, 835–839 (1995).

2. Nagy, L. & Schwabe, J. W. R. Mechanism of the nuclear receptor molecular switch. Trends Biochem Sci 29, 317–324 (2004).

3. Gronemeyer, H., Gustafsson, J.-A. & Laudet, V. Principles for modulation of the nuclear receptor superfamily. Nat Rev Drug Discov 3, 950–964 (2004).

4. Shiau, A. K. et al. The structural basis of estrogen receptor/coactivator recognition and the antagonism of this interaction by tamoxifen. Cell 95, 927–937 (1998).

5. Weikum, E. R., Liu, X. & Ortlund, E. A. The nuclear receptor superfamily: A structural perspective. Protein Sci 27, 1876–1892 (2018).

6. Evans, R. M. The steroid and thyroid hormone receptor superfamily. Science 240, 889–895 (1988).

7. Danielian, P. S., White, R., Lees, J. A. & Parker, M. G. Identification of a conserved region required for hormone dependent transcriptional activation by steroid hormone receptors. The EMBO Journal 11, 1025–1033 (1992).

8. Zhang, C., Zhang, B., Zhang, X., Sun, G. & Sun, X. Targeting Orphan Nuclear Receptors NR4As for Energy Homeostasis and Diabetes. Front. Pharmacol. 11, 587457 (2020).

9. Tata, J. R. Signalling through nuclear receptors. Nat Rev Mol Cell Biol 3, 702–710 (2002).

10. Aranda, A. & Pascual, A. Nuclear hormone receptors and gene expression. Physiol Rev 81, 1269–1304 (2001).

11. A Unified Nomenclature System for the Nuclear Receptor Superfamily. Cell 97, 161–163 (1999).

12. Hopkins, A. L. & Groom, C. R. The druggable genome. Nat Rev Drug Discov 1, 727–730 (2002).

13. Bhasin, M. & Raghava, G. P. S. Classification of Nuclear Receptors Based on Amino Acid Composition and Dipeptide Composition. Journal of Biological Chemistry 279, 23262–23266 (2004).

14. Gao, Y. et al. Using pseudo amino acid composition to predict protein subcellular location: Approached with Lyapunov index, Bessel function, and Chebyshev filter. Amino Acids 28, 373–376 (2005).

15. Wang, P., Xiao, X. & Chou, K.-C. NR-2L: A Two-Level Predictor for Identifying Nuclear Receptor Subfamilies Based on Sequence-Derived Features. PLoS ONE 6, e23505 (2011).

16. Xiao, X., Wang, P. & Chou, K.-C. iNR-PhysChem: A Sequence-Based Predictor for Identifying Nuclear Receptors and Their Subfamilies via Physical-Chemical Property Matrix. PLoS ONE 7, e30869 (2012).

17. NRPred-FS: A Feature Selection based Two-level Predictor for Nuclear Receptors | Abstract. https://www.longdom.org/abstract/nrpredfs-a-feature-selection-based-twolevel-predictor-for-nuclearreceptors-33618.html.

18. Kumar, R., Kumari, B., Srivastava, A. & Kumar, M. NRfamPred: A proteome-scale two level method for prediction of nuclear receptor proteins and their sub-families. Sci Rep 4, 6810 (2014).

19. Keshava Prasad, T. S. et al. Human Protein Reference Database--2009 update. Nucleic Acids Res 37, D767–772 (2009).

20. Ismail, H. D., Saigo, H. & Kc, D. B. RF-NR: Random Forest Based Approach for Improved Classification of Nuclear Receptors. IEEE/ACM Trans Comput Biol Bioinform 15, 1844–1852 (2018).

21. Fischer, A. & Smieško, M. Allosteric Binding Sites On Nuclear Receptors: Focus On Drug Efficacy and Selectivity. Int J Mol Sci 21, E534 (2020).

22. O’Leary, N. A. et al. Reference sequence (RefSeq) database at NCBI: current status, taxonomic expansion, and functional annotation. Nucleic Acids Res 44, D733–745 (2016).

23. Kursa, M. B. & Rudnicki, W. R. Feature Selection with the **Boruta** Package. J. Stat. Soft. 36, (2010).

24. Ding, H. & Li, D. Identification of mitochondrial proteins of malaria parasite using analysis of variance. Amino Acids 47, 329–333 (2015).

25. Mermel, C. H. et al. GISTIC2.0 facilitates sensitive and confident localization of the targets of focal somatic copy-number alteration in human cancers. Genome Biol 12, R41 (2011).

26. Bian, H., Guo, M. & Wang, J. Recognition of Mitochondrial Proteins in Plasmodium Based on the Tripeptide Composition. Front. Cell Dev. Biol. 8, 578901 (2020).

27. Ding, C., Yuan, L.-F., Guo, S.-H., Lin, H. & Chen, W. Identification of mycobacterial membrane proteins and their types using over-represented tripeptide compositions. Journal of Proteomics 77, 321–328 (2012).

28. Yang, L. et al. Identification of Cancerlectins By Using Cascade Linear Discriminant Analysis and Optimal g-gap Tripeptide Composition. CBIO 15, 528–537 (2020).

29. Yang, L., Gao, H., Liu, Z. & Tang, L. Identification of Phage Virion Proteins by Using the g-gap Tripeptide Composition. LOC 16, 332–339 (2019).

30. Wang, K., Li, S., Wang, Q. & Hou, C. Identification of hormone-binding proteins using a novel ensemble classifier. Computing 101, 693–703 (2019).

31. You, Z.-H., Yu, J.-Z., Zhu, L., Li, S. & Wen, Z.-K. A MapReduce based parallel SVM for large-scale predicting protein–protein interactions. Neurocomputing 145, 37–43 (2014).

32. Limongelli, I., Marini, S. & Bellazzi, R. PaPI: pseudo amino acid composition to score human protein-coding variants. BMC Bioinformatics 16, 123 (2015).

33. Chen, Z. et al. Prediction of Ubiquitination Sites by Using the Composition of k-Spaced Amino Acid Pairs. PLoS ONE 6, e22930 (2011).

34. Lv, Z., Wang, D., Ding, H., Zhong, B. & Xu, L. Escherichia Coli DNA N-4-Methycytosine Site Prediction Accuracy Improved by Light Gradient Boosting Machine Feature Selection Technology. IEEE Access 8, 14851–14859 (2020).

35. Li, Y., Zhang, Z., Teng, Z. & Liu, X. PredAmyl-MLP: Prediction of Amyloid Proteins Using Multilayer Perceptron. Computational and Mathematical Methods in Medicine 2020, 1–12 (2020).

36. Xu, R. et al. Identification of DNA-binding proteins by incorporating evolutionary information into pseudo amino acid composition via the top-n-gram approach. Journal of Biomolecular Structure and Dynamics 33, 1720–1730 (2015).

37. Nanni, L., Brahnam, S. & Lumini, A. Wavelet images and Chou’s pseudo amino acid composition for protein classification. Amino Acids 43, 657–665 (2012).

38. Lv, Z., Zhang, J., Ding, H. & Zou, Q. RF-PseU: A Random Forest Predictor for RNA Pseudouridine Sites. Frontiers in Bioengineering and Biotechnology 8, (2020).

39. Sikander, R., Ghulam, A. & Ali, F. XGB-DrugPred: computational prediction of druggable proteins using eXtreme gradient boosting and optimized features set. Sci Rep 12, 5505 (2022).

40. Sequeira, A. M., Lousa, D. & Rocha, M. ProPythia: A Python package for protein classification based on machine and deep learning. Neurocomputing 484, 172–182 (2022).

41. Xiao, X., Shao, S.-H., Huang, Z.-D. & Chou, K.-C. Using pseudo amino acid composition to predict protein structural classes: Approached with complexity measure factor. J. Comput. Chem. 27, 478–482 (2006).

42. Yu, H., Yang, J., Wang, W. & Han, J. Discovering compact and highly discriminative features or feature combinations of drug activities using support vector machines. Proc IEEE Comput Soc Bioinform Conf 2, 220–228 (2003).

43. Lundberg, S. M. & Lee, S.-I. A Unified Approach to Interpreting Model Predictions. in Advances in Neural Information Processing Systems vol. 30 (Curran Associates, Inc., 2017).

44. Shapley, L. S. 17. A Value for n-Person Games. in Contributions to the Theory of Games (AM-28), Volume II (eds. Kuhn, H. W. & Tucker, A. W.) 307–318 (Princeton University Press, 1953). doi:10.1515/9781400881970-018.

45. Neduva, V. et al. Systematic Discovery of New Recognition Peptides Mediating Protein Interaction Networks. PLoS Biol 3, e405 (2005).

46. Ning, Q., Qi, Z., Wang, Y., Deng, A. & Chen, C. FCCCSR_Glu: a semi-supervised learning model based on FCCCSR algorithm for prediction of glutarylation sites. Briefings in Bioinformatics bbac421 (2022) doi:10.1093/bib/bbac421.

47. Li, S. et al. pCysMod: Prediction of Multiple Cysteine Modifications Based on Deep Learning Framework. Front. Cell Dev. Biol. 9, 617366 (2021).

48. Vroling, B. et al. NucleaRDB: information system for nuclear receptors. Nucleic Acids Research 40, D377–D380 (2012).

49. https://rescued.omnipathdb.org/.

50. Chen, Z. et al. iFeature: a Python package and web server for features extraction and selection from protein and peptide sequences. Bioinformatics 34, 2499–2502 (2018).

51. Komer, B., Bergstra, J. & Eliasmith, C. Hyperopt-Sklearn: Automatic Hyperparameter Configuration for Scikit-Learn. in 32–37 (2014). doi:10.25080/Majora-14bd3278-006.

