## Supplementary information for "NRPreTo: A Machine Learning Based Nuclear Receptor and Subfamily Prediction Tool"

### NRPreTo : A Two-level Machine Learning Based Nuclear Receptor Subfamily Prediction using Protein Feature Spectrum

**Table S1.** Description of the subfamily names and their members across the two benchmark training datasets

| **Subfamily** | **protein name** | **BD1** | | **BD2** | |
|---|---|---|---|---|---|
| | | Training | Independent | Training | Independent |
| NR0 | DAX-like and Knirps-like (DAX and SHP), knirps related proteins | 48 | 45 | 50 | 231 |
| NR1 | Thyroid hormone-like, retinoic acid, peroxisome proliferators activated, Vitamin D3-like, RAR related orphan receptors | 82 | 1090 | 36 | 127 |
| NR2 | Hepatocyte nuclear factor 4-like, tailless-like, Retinoic acid receptor (RXR), COUP proteins | 68 | 668 | 37 | 148 |
| NR3 | Estrogen-like, glucocorticoid-like | 33 | 671 | 7 | 23 |
| NR4 | Nerve growth factor I-B-like | 11 | 108 | 12 | 33 |
| NR5 | Fushi tarazu-F1-like | 15 | 136 | 5 | 0 |
| NR6 | Germ cell-like | 10 | 31 | 12 | 6 |
| Non-NR | Non-Nuclear Receptor | 1000 | 1064 | 500 | 500 |
| | **TOTAL** | **1267** | **3813** | **659** | **1067** |

**Table S2.** Sequences predicted as Non-NR by NrPreTo but as NR by NRfamPred and their comparison with HRPD and UNIPROT predictions

| **Refseq** | **Name** | **NrPreTo** | **NRfamPred** | **HRPD** | **UNIPROT** |
|---|---|---|---|---|---|
| NP_004917.2 | Butyrate response factor 1 | NonNR | 1 | NonNR | NonNR |
| NP_005420.1 | Vascular endothelial growth factor C | NonNR | 1 | NonNR | NonNR |
| NP_001120968.1 | Protein sprouty homolog 4 isoform 2 | NonNR | 3 | NonNR | NonNR |
| NP_004548.3 | Neurogenic locus notch homolog protein 4 preproprotein | NonNR | 3 | NonNR | NonNR |
| NP_005928.2 | Protein AF-17 | NonNR | 3 | NonNR | NonNR |
| NP_076421.2 | Protein delta homolog 2 isoform a precursor | NonNR | 3 | NonNR | NonNR |
| NP_112226.2 | Protein sprouty homolog 4 isoform 1 | NonNR | 3 | NonNR | NonNR |
| NP_113621.1 | Membrane frizzled-related protein | NonNR | 3 | NonNR | NonNR |
| NP_775960.4 | Protein crumbs homolog 2 precursor | NonNR | 3 | NonNR | NonNR |
| NP_996262.1 | Protein delta homolog 2 isoform a precursor | NonNR | 3 | NonNR | NonNR |
| NP_008933.2 | Proteasomal ubiquitin receptor ADRM1 isoform 1 | NonNR | 5 | NonNR | NonNR |
| NP_783163.1 | Proteasomal ubiquitin receptor ADRM1 isoform 1 | NonNR | 5 | NonNR | NonNR |

**Table S3.** Sequences with predictions in NrPreTo but not by NRfamPred and comparison with HPRD and UNIPROT

| S.No | Refseq | Name | NrPreTo | NRfamPred | HRPD | UNIPROT |
|---|---|---|---|---|---|---|
| 1 | NP_000452.2 | Thyroid hormone receptor, beta | 1 | Unreported | 1 | 1 |
| 2 | NP_001017535.1 | Vitamin D receptor | 1 | Unreported | 1 | 1 |
| 3 | NP_001121648.1 | Thyroid hormone receptor, beta | 1 | Unreported | 1 | 1 |
| 4 | NP_001121649.1 | Thyroid hormone receptor, beta | 1 | Unreported | 1 | 1 |
| 5 | NP_001138774.1 | Retinoic acid receptor alpha | 1 | Unreported | 1 | 1 |
| 6 | NP_002934.1 | RAR related orphan receptor A | 1 | Unreported | 1 | 1 |
| 7 | NP_008845.2 | RAR related orphan receptor B | 1 | Unreported | 1 | 1 |
| 8 | NP_056953.2 | Peroxisome proliferator activated receptor gamma | 1 | Unreported | 1 | 1 |
| 9 | NP_057236.1 | Retinoic acid receptor beta | 1 | Unreported | 1 | 1 |
| 10 | NP_599022.1 | RAR related orphan receptor A | 1 | Unreported | 1 | 1 |
| 11 | NP_599023.1 | RAR related orphan receptor A | 1 | Unreported | 1 | 1 |
| 12 | NP_599024.1 | RAR related orphan receptor A | 1 | Unreported | 1 | 1 |
| 13 | NP_619725.2 | Peroxisome proliferator activated receptor gamma | 1 | Unreported | 1 | 1 |
| 14 | NP_619726.2 | Peroxisome proliferator activated receptor gamma | 1 | Unreported | 1 | 1 |
| 15 | NP_955366.1 | Thyroid hormone receptor alpha | 1 | Unreported | 1 | 1 |
| 16 | NP_005028.4 | Peroxisome proliferator activated receptor gamma | 1 | Unreported | 1 | 1 |
| 17 | NP_008848.1 | Retinoid X receptor, gamma | 2 | Unreported | 2 | 2 |
| 18 | NP_001011645.1 | Androgen receptor | 3 | Unreported | 3 | 3 |
| 19 | NP_004442.3 | Estrogen-related receptor alpha | 3 | Unreported | 3 | 3 |
| 20 | NP_000466.2 | Orphan nuclear receptor DAX1 | 0 | Unreported | 0 | 0 |
| 21 | NP_068804.1 | Orphan nuclear receptor SHP | 0 | Unreported | 0 | 0 |
| 22 | NP_001070937.1 | Nuclear receptor subfamily 1 group I member 3 isoform 6 | 1 | Unreported | 1 | 1 |
| 23 | NP_001070938.1 | Nuclear receptor subfamily 1 group I member 3 isoform 11 | 1 | Unreported | 1 | 1 |
| 24 | NP_001070939.1 | Nuclear receptor subfamily 1 group I member 3 isoform 5 | 1 | Unreported | 1 | 1 |
| 25 | NP_001070940.1 | Nuclear receptor subfamily 1 group I member 3 isoform 9 | 1 | Unreported | 1 | 1 |
| 26 | NP_001070941.1 | Nuclear receptor subfamily 1 group I member 3 isoform 12 | 1 | Unreported | 1 | 1 |
| 27 | NP_001070942.1 | Nuclear receptor subfamily 1 group I member 3 isoform 8 | 1 | Unreported | 1 | 1 |
| 28 | NP_001070943.1 | Nuclear receptor subfamily 1 group I member 3 isoform 15 | 1 | Unreported | 1 | 1 |
| 29 | NP_001070944.1 | Nuclear receptor subfamily 1 group I member 3 isoform 13 | 1 | Unreported | 1 | 1 |
| 30 | NP_001070945.1 | Nuclear receptor subfamily 1 group I member 3 isoform 14 | 1 | Unreported | 1 | 1 |
| 31 | NP_001070946.1 | Nuclear receptor subfamily 1 group I member 3 isoform 7 | 1 | Unreported | 1 | 1 |
| 32 | NP_001070947.1 | Nuclear receptor subfamily 1 group I member 3 isoform 10 | 1 | Unreported | 1 | 1 |
| 33 | NP_001070948.1 | Nuclear receptor subfamily 1 group I member 3 | 1 | Unreported | 1 | 1 |
| 34 | NP_001070949.1 | Nuclear receptor subfamily 1 group I member 3 isoform 4 | 1 | Unreported | 1 | 1 |
| 35 | NP_001070950.1 | Nuclear receptor subfamily 1 group I member 3 isoform 1 | 1 | Unreported | 1 | 1 |
| 36 | NP_001123573.1 | Nuclear receptor subfamily 1, group H member 3 | 1 | Unreported | 1 | 1 |
| 37 | NP_001123574.1 | Nuclear receptor subfamily 1, group H member 3 | 1 | Unreported | 1 | 1 |
| 38 | NP_001138897.1 | Nuclear receptor subfamily 1, group D, member 2 | 1 | Unreported | 1 | 1 |
| 39 | NP_005113.1 | Nuclear receptor subfamily 1 group I member 3 isoform 3 | 1 | Unreported | 1 | 1 |
| 40 | NP_005117.3 | Nuclear receptor subfamily 1, group D, member 2 | 1 | Unreported | 1 | 1 |
| 41 | NP_148934.1 | Nuclear receptor subfamily 1, group I, member 2 | 1 | Unreported | 1 | 1 |
| 42 | NP_001027458.1 | Nuclear receptor subfamily 2 group C | 2 | Unreported | 2 | 2 |
| 43 | NP_001120834.1 | Nuclear receptor subfamily 2 group C member 1 isoform c | 2 | Unreported | 2 | 2 |
| 44 | NP_001138627.1 | Nuclear receptor subfamily 2, group F, member 2 | 2 | Unreported | 2 | 2 |
| 45 | NP_001138628.1 | Nuclear receptor subfamily 2, group F, member 2 | 2 | Unreported | 2 | 2 |
| 46 | NP_001138629.1 | Nuclear receptor subfamily 2, group F, member 2 | 2 | Unreported | 2 | 2 |
| 47 | NP_003260.1 | Nuclear receptor TLX | 2 | Unreported | 2 | 2 |
| 48 | NP_003288.2 | Nuclear receptor subfamily 2 group C member 1 isoform a | 2 | Unreported | 2 | 2 |
| 49 | NP_003289.2 | Nuclear receptor subfamily 2 group C member 2 isoform 1 | 2 | Unreported | 2 | 2 |
| 50 | NP_000167.1 | Glucocorticoid receptor | 3 | Unreported | 3 | 3 |
| 52 | NP_001018084.1 | Glucocorticoid receptor | 3 | Unreported | 3 | 3 |
| 53 | NP_001018085.1 | Glucocorticoid receptor | 3 | Unreported | 3 | 3 |
| 54 | NP_001018086.1 | Glucocorticoid receptor | 3 | Unreported | 3 | 3 |

| No. | Refseq | Name | NrPreTo | NRfamPred | HRPD | UNIPROT |
|---|---|---|---|---|---|---|
| 55 | NP_001018087.1 | Glucocorticoid receptor | 3 | Unreported | 3 | 3 |
| 56 | NP_001018661.1 | Glucocorticoid receptor | 3 | Unreported | 3 | 3 |
| 57 | NP_001019265.1 | Glucocorticoid receptor | 3 | Unreported | 3 | 3 |
| 58 | NP_008912.2 | Nuclear receptor subfamily 4, group A, member 3 | 4 | Unreported | 4 | 4 |
| 59 | NP_775291.1 | Nuclear receptor subfamily 4, group A, member 3 | 4 | Unreported | 4 | 4 |
| 60 | NP_775292.1 | Nuclear receptor subfamily 4, group A, member 3 | 5 | Unreported | 5 | 5 |
| 61 | NP_995582.1 | Nuclear receptor subfamily 5, group A, member 2 | 5 | Unreported | 5 | 5 |

Unreported refers to sequences which have no reported predictions in NrfamPred

**Table S4.** Sequences predicted by NrPreTo having annotations in UNIPROT but not in NRfamPred and HPRD

| S.No | Refseq | Name | NrPreTo | NRfamPred | HRPD | UNIPROT |
|---|---|---|---|---|---|---|
| 1 | NP_001017536.1 | Vitamin D3 receptor | 1 | Unreported | NID | 1 |
| 2 | NP_001165289.1 | Peroxisome proliferator-activated receptor delta | 1 | Unreported | NID | 1 |
| 3 | NP_001165290.1 | Peroxisome proliferator-activated receptor delta | 1 | Unreported | NID | 1 |
| 4 | NP_001177847.1 | Thyroid hormone receptor alpha | 1 | Unreported | NID | 1 |
| 5 | NP_001177848.1 | Thyroid hormone receptor alpha | 1 | Unreported | NID | 1 |
| 6 | NP_001193906.1 | Bile acid receptor (Farnesoid X-activated receptor) | 1 | Unreported | NID | 1 |
| 7 | NP_001193908.1 | Bile acid receptor (Farnesoid X-activated receptor) | 1 | Unreported | NID | 1 |
| 8 | NP_001193921.1 | Bile acid receptor (Farnesoid X-activated receptor) | 1 | Unreported | NID | 1 |
| 9 | NP_001193922.1 | Bile acid receptor (Farnesoid X-activated receptor) | 1 | Unreported | NID | 1 |
| 10 | NP_001230659.1 | Retinoic acid receptor gamma, RAR-gamma | 1 | Unreported | NID | 1 |
| 11 | NP_001230661.1 | Retinoic acid receptor gamma, RAR-gamma | 1 | Unreported | NID | 1 |
| 12 | NP_001239563.1 | Thyroid hormone receptor beta | 1 | Unreported | NID | 1 |
| 13 | NP_001277145.1 | Retinoic acid receptor beta, RAR-beta (HBV-activated protein) | 1 | Unreported | NID | 1 |
| 14 | NP_001277195.1 | Retinoic acid receptor beta, RAR-beta (HBV-activated protein) | 1 | Unreported | NID | 1 |
| 15 | NP_001277229.1 | Retinoic acid receptor beta variant 1 | 1 | Unreported | NID | 1 |
| 16 | NP_001243500.1 | Retinoic acid receptor RXR-gamma | 2 | Unreported | NID | 2 |
| 17 | NP_001245284.1 | Hepatocyte nuclear factor 4-alpha | 2 | Unreported | NID | 2 |
| 18 | NP_001257330.1 | Retinoic acid receptor RXR-beta | 2 | Unreported | NID | 2 |
| 19 | NP_001273031.1 | Nuclear receptor subfamily 2 group E member 1 | 2 | Unreported | NID | 2 |
| 20 | NP_001274111.1 | Hepatocyte nuclear factor 4-alpha, HNF-4-alpha | 2 | Unreported | NID | 2 |
| 21 | NP_001274112.1 | Hepatocyte nuclear factor 4-alpha, HNF-4-alpha | 2 | Unreported | NID | 2 |
| 22 | NP_001274113.1 | Hepatocyte nuclear factor 4-alpha | 2 | Unreported | NID | 2 |
| 23 | NP_001278623.1 | Nuclear receptor subfamily 2 group C member 2 | 2 | Unreported | NID | 2 |
| 24 | NP_001278850.1 | Retinoic acid receptor RXR-alpha | 2 | Unreported | NID | 2 |
| 25 | NP_001317490.1 | Hepatocyte nuclear factor 4-gamma | 2 | Unreported | NID | 2 |
| 26 | NP_001159576.1 | Mineralocorticoid receptor | 3 | Unreported | NID | 3 |
| 27 | NP_001189403.1 | Progesterone receptor, PR | 3 | Unreported | NID | 3 |
| 28 | NP_001191187.1 | Glucocorticoid receptor, GR | 3 | Unreported | NID | 3 |
| 29 | NP_001191188.1 | Glucocorticoid receptor, GR | 3 | Unreported | NID | 3 |
| 30 | NP_001191189.1 | Glucocorticoid receptor, GR | 3 | Unreported | NID | 3 |
| 31 | NP_001191190.1 | Glucocorticoid receptor, GR | 3 | Unreported | NID | 3 |
| 32 | NP_001191191.1 | Glucocorticoid receptor, GR | 3 | Unreported | NID | 3 |
| 33 | NP_001191192.1 | Glucocorticoid receptor, GR | 3 | Unreported | NID | 3 |
| 34 | NP_001191193.1 | Glucocorticoid receptor, GR | 3 | Unreported | NID | 3 |
| 35 | NP_001201831.1 | Estrogen receptor beta | 3 | Unreported | NID | 3 |
| 36 | NP_001230438.1 | Estrogen-related receptor gamma | 3 | Unreported | NID | 3 |
| 37 | NP_001230439.1 | Estrogen-related receptor gamma | 3 | Unreported | NID | 3 |
| 38 | NP_001230440.1 | Estrogen-related receptor gamma | 3 | Unreported | NID | 3 |
| 39 | NP_001230441.1 | Estrogen-related receptor gamma | 3 | Unreported | NID | 3 |
| 40 | NP_001230442.1 | Estrogen-related receptor gamma | 3 | Unreported | NID | 3 |
| 41 | NP_001230443.1 | Estrogen-related receptor gamma | 3 | Unreported | NID | 3 |
| 42 | NP_001230444.1 | Estrogen-related receptor gamma | 3 | Unreported | NID | 3 |
| 43 | NP_001230447.1 | Estrogen-related receptor gamma | 3 | Unreported | NID | 3 |
| 44 | NP_001230448.1 | Estrogen-related receptor gamma | 3 | Unreported | NID | 3 |
| 45 | NP_001258090.1 | Progesterone receptor, PR | 3 | Unreported | NID | 3 |
| 46 | NP_001258805.1 | Estrogen receptor beta, ER-beta | 3 | Unreported | NID | 3 |
| 47 | NP_001258806.1 | Estrogen receptor beta, ER-beta | 3 | Unreported | NID | 3 |
| 48 | NP_001269379.1 | Steroid hormone receptor ERR1 | 3 | Unreported | NID | 3 |
| 49 | NP_001269380.1 | Steroid hormone receptor ERR1 | 3 | Unreported | NID | 3 |
| 50 | NP_001278159.1 | Estrogen receptor | 3 | Unreported | NID | 3 |

| S.No | Refseq | Name | NrPreTo | NRfamPred | HRPD | UNIPROT |
|---|---|---|---|---|---|---|
| 51 | NP_001278170.1 | Estrogen receptor | 3 | Unreported | NID | 3 |
| 52 | NP_001278641.1 | Estrogen receptor | 3 | Unreported | NID | 3 |
| 53 | NP_001278652.1 | Estrogen receptor | 3 | Unreported | NID | 3 |
| 54 | NP_001315029.1 | Estrogen receptor | 3 | Unreported | NID | 3 |
| 55 | NP_001334990.1 | Androgen receptor | 3 | Unreported | NID | 3 |
| 56 | NP_001334992.1 | Androgen receptor | 3 | Unreported | NID | 3 |
| 57 | NP_001189162.1 | Nuclear receptor subfamily 4 group A member Testicular receptor 3 | 4 | Unreported | NID | 4 |
| 58 | NP_001189163.1 | Nuclear receptor subfamily 4 group A member 1 | 4 | Unreported | NID | 4 |
| 59 | NP_001263393.1 | (Alpha-1-fetoprotein transcription factor) | 5 | Unreported | NID | 5 |
| 60 | NP_001265475.1 | Nuclear receptor subfamily 6 group A member 1 (Germ cell nuclear factor, GCNF, hGCNF) | 6 | Unreported | NID | 6 |

NID refers to class not reported in database

**Table S5.** Novel sequences predicted by NrPreTo not reported in UNIPROT, NRfamPred and HPRD

| S.No | Refseq | Name | NrPreTo | NRfamPred | HRPD | UNIPROT |
|---|---|---|---|---|---|---|
| 1 | NP_001337051.1 | Estrogen-related-receptor-gamma-isoform-2 | 3 | Unreported | NID | NID |
| 2 | NP_001337052.1 | Estrogen-related-receptor-gamma-isoform-2 | 3 | Unreported | NID | NID |
| 3 | NP_001337053.1 | Estrogen-related-receptor-gamma-isoform-2 | 3 | Unreported | NID | NID |
| 4 | NP_001337054.1 | Estrogen-related-receptor-gamma-isoform-2 | 3 | Unreported | NID | NID |
| 5 | NP_001341595.2 | Peroxisome-proliferator-activated-receptor gamma-isoform-1 | 1 | Unreported | NID | NID |
| 6 | NP_001341596.2 | Peroxisome-proliferator-activated-receptor gamma-isoform-1 | 1 | Unreported | NID | NID |
| 7 | NP_001341597.1 | Peroxisome-proliferator-activated-receptor gamma-isoform-4 | 1 | Unreported | NID | NID |
| 8 | NP_001341599.1 | Peroxisome-proliferator-activated-receptor gamma-isoform-6 | 1 | Unreported | NID | NID |
| 9 | NP_001341637.1 | Thyroid-hormone-receptor-beta-isoform-a | 1 | Unreported | NID | NID |
| 10 | NP_001341638.1 | Thyroid-hormone-receptor-beta-isoform-a | 1 | Unreported | NID | NID |
| 11 | NP_001341639.1 | Thyroid-hormone-receptor-beta-isoform-a | 1 | Unreported | NID | NID |
| 12 | NP_001341640.1 | Thyroid-hormone-receptor-beta-isoform-a | 1 | Unreported | NID | NID |
| 13 | NP_001341641.1 | Thyroid-hormone-receptor-beta-isoform-a | 1 | Unreported | NID | NID |
| 14 | NP_001341642.1 | Thyroid-hormone-receptor-beta-isoform-a | 1 | Unreported | NID | NID |
| 15 | NP_001341643.1 | Thyroid-hormone-receptor-beta-isoform-b | 1 | Unreported | NID | NID |
| 16 | NP_001341644.1 | Thyroid-hormone-receptor-beta-isoform-b | 1 | Unreported | NID | NID |
| 17 | NP_001341748.1 | Mineralocorticoid-receptor-isoform-2 | 3 | Unreported | NID | NID |
| 18 | NP_001349801.1 | Peroxisome-proliferator-activated-receptor alpha-isoform-1 | 1 | Unreported | NID | NID |
| 19 | NP_001349802.1 | Peroxisome-proliferator-activated-receptor alpha-isoform-1 | 1 | Unreported | NID | NID |
| 20 | NP_001350524.1 | Oxysterols-receptor-LXR-alpha-isoform-5 | 1 | Unreported | NID | NID |
| 21 | NP_001351014.1 | Vitamin-D3-receptor-isoform-VDRA | 1 | Unreported | NID | NID |
| 22 | NP_001351109.1 | Glucocorticoid-receptor-isoform-alpha | 3 | Unreported | NID | NID |
| 23 | NP_001351110.1 | Glucocorticoid-receptor-isoform-alpha | 3 | Unreported | NID | NID |
| 24 | NP_001351111.1 | Glucocorticoid-receptor-isoform-alpha | 3 | Unreported | NID | NID |
| 25 | NP_001351112.1 | Glucocorticoid-receptor-isoform-gamma | 3 | Unreported | NID | NID |
| 26 | NP_001351113.1 | Glucocorticoid-receptor-isoform-gamma | 3 | Unreported | NID | NID |
| 27 | NP_001351114.1 | Glucocorticoid-receptor-isoform-gamma | 3 | Unreported | NID | NID |
| 28 | NP_001351952.1 | Nuclear-receptor-ROR-beta-isoform-2 | 1 | Unreported | NID | NID |
| 29 | NP_001361190.2 | Peroxisome-proliferator-activated-receptor gamma-isoform-3 | 1 | Unreported | NID | NID |
| 30 | NP_001361191.2 | Peroxisome-proliferator-activated-receptor gamma-isoform-3 | 1 | Unreported | NID | NID |
| 31 | NP_001361192.2 | Peroxisome-proliferator-activated-receptor gamma-isoform-1 | 1 | Unreported | NID | NID |
| 32 | NP_001361193.2 | Peroxisome-proliferator-activated-receptor gamma-isoform-1 | 1 | Unreported | NID | NID |
| 33 | NP_001361194.1 | Peroxisome-proliferator-activated-receptor gamma-isoform-7 | 1 | Unreported | NID | NID |
| 34 | NP_001361195.1 | Peroxisome-proliferator-activated-receptor gamma-isoform-8 | 1 | Unreported | NID | NID |
| 35 | NP_001361590.1 | Vitamin-D3-receptor-isoform-VDRA | 1 | Unreported | NID | NID |
| 36 | NP_001361591.1 | Vitamin-D3-receptor-isoform-VDRA | 1 | Unreported | NID | NID |
| 37 | NP_001361751.1 | Thyroid-hormone-receptor-beta-isoform-a | 1 | Unreported | NID | NID |
| 38 | NP_001361752.1 | Thyroid-hormone-receptor-beta-isoform-a | 1 | Unreported | NID | NID |
| 39 | NP_001361753.1 | Thyroid-hormone-receptor-beta-isoform-a | 1 | Unreported | NID | NID |
| 40 | NP_001361754.1 | Thyroid-hormone-receptor-beta-isoform-a | 1 | Unreported | NID | NID |
| 41 | NP_001361755.1 | Thyroid-hormone-receptor-beta-isoform-a | 1 | Unreported | NID | NID |
| 42 | NP_001361756.1 | Thyroid-hormone-receptor-beta-isoform-c | 1 | Unreported | NID | NID |
| 43 | NP_001366109.1 | Steroid-hormone-receptor-ERR2-isoform-2 | 3 | Unreported | NID | NID |
| 44 | NP_001372497.1 | Estrogen-receptor-isoform-1 | 3 | Unreported | NID | NID |
| 45 | NP_001372498.1 | Estrogen-receptor-isoform-1 | 3 | Unreported | NID | NID |
| 46 | NP_001372499.1 | Estrogen-receptor-isoform-5 | 3 | Unreported | NID | NID |
| 47 | NP_001372500.1 | Estrogen-receptor-isoform-5 | 3 | Unreported | NID | NID |
| 48 | NP_001372501.1 | Estrogen-receptor-isoform-5 | 3 | Unreported | NID | NID |
| 49 | NP_001380870.1 | Peroxisome-proliferator-activated-receptor-alpha-isoform-1 | 1 | Unreported | NID | NID |
| 50 | NP_001380871.1 | Peroxisome-proliferator-activated-receptor-alpha-isoform-1 | 1 | Unreported | NID | NID |
| 51 | NP_001380872.1 | Peroxisome-proliferator-activated-receptor-alpha-isoform-1 | 1 | Unreported | NID | NID |
| 52 | NP_001380873.1 | Peroxisome-proliferator-activated-receptor-alpha-isoform-1 | 1 | Unreported | NID | NID |
| 53 | NP_001380874.1 | Peroxisome-proliferator-activated-receptor-alpha-isoform-1 | 1 | Unreported | NID | NID |

| S.No | Refseq | Name | NrPreTo | NRfamPred | HRPD | UNIPROT |
|---|---|---|---|---|---|---|
| 54 | NP_005028.5 | Peroxisome-proliferator-activated-receptor-gamma-isoform-1 | 1 | Unreported | NID | NID |
| 55 | NP_619725.3 | Peroxisome-proliferator-activated-receptor-gamma-isoform-1 | 1 | Unreported | NID | NID |
| 56 | NP_619726.3 | Peroxisome-proliferator-activated-receptor-gamma-isoform-1 | 1 | Unreported | NID | NID |
| 57 | NP_001191194.1 | Glucocorticoid receptor isoform GR-P | 3 | Unreported | NID | NID |
| 58 | NP_001277206.1 | Retinoic acid receptor beta isoform 5 | 1 | Unreported | NID | NID |
| 59 | NP_001317544.2 | Peroxisome proliferator-activated receptor gamma isoform 3 | 1 | Unreported | NID | NID |

NID refers to class not in database

**Table S6**. Level 1 prediction performance of "BD1 train only" and "BD2 "train only" RF models on external datasets over 15 runs

| **External Datasets** | **BD1 train only \| BD2 train only** | | | |
|---|---|---|---|---|
| | **Acc** | **F1** | **MCC** | **AUC** |
| HPRD | 0.9890 ± 0.0001 \| 0.9894 ± 0.0002 | 0.9312 ± 0.0061 \| 0.9537 ± 0.0029 | 0.8726 ± 0.0121 \| 0.9178 ± 0.0059 | 0.9341 ± 0.0016 \| 0.9415 ± 0.0001 |
| Refseq | 0.9892 ± 0.0001 \| 0.9896 ± 0.0001 | 0.9423 ± 0.0056 \| 0.9609 ± 0.0033 | 0.8947 ± 0.0113 \| 0.9323 ± 0.0068 | 0.9412 ± 0.0013 \| 0.9456 ± 0.0005 |

On each row values separated by '|'means prediction performance of a Random Forest-model trained on BD1 | BD2 and tested on external dataset over 15 runs to predict NR vs. Non-NR for human proteins on HPRD and RefSeq databases. Acc: accuracy. MCC: Matthew's Correlation Coefficient. AUC: Area under the curve.

**Table S7.** Level 2 prediction performance of RF BD1 combined model and BD2 combined model (combination of train and independent dataset) on external datasets over 15 runs.

| External Datasets | Combined BD1 \| Combined BD2 | | | |
|---|---|---|---|---|
| | **Acc** | **F1** | **MCC** | **AUC** |
| HPRD | 0.9809 ± 0.0011 \| 0.9811 ± 0.0021 | 0.9574 ± 0.0018 \| 0.9621 ± 0.0021 | 0.9771 ± 0.0001 \| 0.9776 ± 0.0015 | 0.9560 ± 0.0065 \| 0.9630 ± 0.0077 |
| Refseq | 0.9722 ± 0.0001 \| 0.9724 ± 0.0002 | 0.9455 ± 0.0023 \| 0.9519 ± 0.0031 | 0.9632 ± 0.0002 \| 0.9637 ± 0.0020 | 0.9522 ± 0.0013 \| 0.9642 ± 0.0091 |

On each row values separated by '|'means prediction performance of a Random Forest -model trained on BD1 | BD2 and tested on external dataset over 15 runs to predict NR sub-class proteins on HPRD and RefSeq databases. Acc: accuracy. MCC: Matthew's Correlation Coefficient. AUC: Area under the curve.

**Table S8.** Summary of the different hyperparameters and their ranges tuned in the study

| Model | Hyperparameters : Range |
|---|---|
| RF | n_estimators: uniform_distribution [75,125]<br>max_samples: uniform_distribution[0,1]<br>max_depth: uniform_distribution[1,100]<br>min_samples_split : uniform_distribution[2,10]<br>min_samples_leaf: uniform_distribution[1, 10] |
